# The deep winding at the brain surface: replicating a historical report associating the ‘bridged’ central sulcus with the pli de passage fronto-pariétal moyen

**DOI:** 10.1101/2024.12.30.630789

**Authors:** R. Schweizer, A. M. Muellen, J. Stropel

## Abstract

In 1876, the anatomist Heschl surveyed 1,087 brains identifying six cases of a unilateral ‘bridged’ central sulcus (CS) at the brain surface. He also measured the height of a minor ‘deep winding’ at the same location within the CS in the remaining 1,081 brains, reporting a distribution skewed towards significantly increased heights. These observations supported his hypothesis that the ‘bridged’ CS represents an extreme form of the ‘deep winding’ within the CS. In this replication we examined structural MRI data from an equally large dataset of 1,112 participants of the Human Connectome Project young adult (HCP-YA) dataset. Through visual inspection, we identified nine cases of a ‘bridged’ CS, confirming its prevalence of less than 1%. The height of the ‘deep winding’, referred to in the HCP-YA dataset as the pli de passage fronto-pariétal moyen (PPfpm), was extracted from 1,983 MRI-based hemispheric depth profiles. The resulting PPfpm height distribution, although wider, still mirrored Heschl’s findings, showing a similar skew towards larger heights. Further analyses of the twin data within the HCP-YA dataset indicated a slightly increased prevalence of the ‘bridged’ CS in monozygotic and dizygotic twins compared to non-twin individuals, though no concordance of ‘bridged’ CS was observed in monozygotic twin pairs. This replication study validates both of Heschl’s observations, describes additional factors that might influence the prevalence of the ‘bridged’ CS, and refines the characterization of the ‘deep winding’ height distribution. Together, these findings reaffirm and expand historical insights into the intricate anatomical organization of the CS.

## Introduction

Why replicating a historical anatomical study from 1877? This study resolves an anatomical ambiguity by linking a single case description of a ‘bridged’ central sulcus (CS) (Wagner 1862) with the common ‘deep winding’ observed at the same location at the fundus of the CS (Heschl 1877). Since Heschl’s study is the only one addressing the relationship between the ‘bridged’ CS and the ‘deep winding’, and uniquely employed a statistical approach with a very large dataset, the present replication sought to determine if Heschl’s assumptions could be confirmed. Additionally, by applying modern analysis methods, our second aim was to generate a refined modern dataset of ‘bridged’ central sulci and the distribution of ‘deep winding’ heights, providing deeper insights into the detailed structural layout of the CS.

In 1862, Wagner described a particular finding in a single brain specimen: “bridges” across the CS, characterized as a prominent connection between the precentral and postcentral gyrus at the upper third of the left hemispheric CS. This finding gained recognition through a popular brain anatomy book, which stated that the CS is “never or extremely rarely being bridged by a secondary winding” (Ecker 1869; translation by the authors). Over the following 50 years, a limited number of case reports describing a ‘bridged’ CS were published (von Monakow 1905, for an overview see Schweizer et al. 2014a), though reports with contradictory results also appeared, with some explicitly noting they have never observed this anatomical variation (Turner, Bischoff, reported in Ecker 1869, Broca 1888, Retzius 1896). Broca’s account is particularly notable as he indirectly linked the ‘bridged’ CS with a deep annectant gyrus within the CS: the “pli de passage fronto-pariétal moyen” (Broca 1888) with “pli de passage” implying a traversing gyrus between lobes (Gratiolet 1854 cited in Broca 1888), “fronto-pariétal”, indicating a position between fontal and parietal lobes, and “moyen” signifying a position in the middle along the CS. This term remains in use today (Alkhadi and Kollias 2004, Boling et al. 1999, 2004), partly due to the absence of a modern anatomical nomenclature. Broca emphasizes that the pli de passage fronto-pariétal moyen (PPfpm) lies deep within the CS, “never approaches the surface” (translation by the authors) and that he had only observed a superficial variation once, in a brain with numerous severe anomalies. Despite the continued relevance of Broca’s comprehensive anatomical descriptions, his claim that the PPfpm is always a deep structure at the fundus of the CS had already been challenged 11 years earlier by Heschl’s 1877 publication.

In a short note on the cerebral cortex ‘deep windings’ and the ‘bridged’ CS, Heschl reported six cases of a unilateral ‘bridged’ CS in a dataset of 1087 brain specimens. He also detailed the height distribution of the ‘deep winding’ at the same CS location in the remaining 1081 brains, identifying 142 cases where ‘deep windings’ rose significantly beyond the typical height, some nearly reaching the brain’s surface. These findings strongly supported his proposition that the ‘bridged’ CS represents a variant of the common ‘deep winding’ at the same location in the fundus of the CS. We came across Heschl’s historical report during our investigation of the historical brain specimen of Wagner’s original description, which was serendipitously discovered as the mix up with the brain of C.F. Gauss (Schweizer et al. 2014b). Through MR images we demonstrated that the “bridges” are fully formed gyri connecting the pre- and postcentral gyri from the fundus to the brain surface, resulting in a divided, discontinuous CS (Schweizer et al. 2014a).

Heschl’s detailed anatomical observation inspired us to replicate (Block et al. 2018, Brandt et al. 2014) his findings using structural whole-brain MR images of the 1200 subjects from the Human Connectome Project young adult (HCP-YA) data set (Ref. HCP-YA). We employed two distinct methodological approaches to assess the prevalence and the height distribution. The ‘bridged’ central sulci were identified through visual inspections of MRI surface reconstructions (Schweizer et al. 2018a, Schweizer et al. 2018b), presumably similar to Heschl’s historical approach, thus ensuring that any discrepancies in prevalence could be attributed to differences in the two dataset populations rather than in the methodology. The height of the deep annectant gyrus within the CS was estimated through an automized detection of landmarks (Müllen & Schweizer, 2022) extracted from the MRI-based CS depth profiles (Cykowski et al. 2008). Differences in the height distribution are therefore likely due to methodological differences. To acknowledge the two historical terms used for the deep annectant gyrus within the CS we use Heschl’s nomenclature “deep winding” related to the historical dataset and Broca’s term ‘PPfpm’ related to the HCP-YA dataset.

The high percentage of monozygotic and dizygotic twin pairs in the HCP-YA dataset enabled an extension to the replication: an exploration of the prevalence of the ‘bridged’ CS in twin individuals and the potential increase in prevalence of such cases in monozygotic twin pairs. This twin dataset provided a unique opportunity to further investigate the genetic and environmental influences on the occurrence of the ‘bridged CS, particularly by comparing its prevalence within monozygotic twin pairs and between monozygotic and dizygotic twins and non-twin siblings.

## Material and Methods

### MRI Data

We analyzed 1112 unprocessed whole brain structural MRI datasets (3T, T1-weighted MPRAGE, spatial resolution: 0.7 x 0.7 x 0.7mm^3^) obtained from the Human Connectome Project 1200 young adult subject release (HCP-YA; S1200 Release, February 2017) (van Essen et al. 2012). Additional information on participant’s age, gender, self-reported family affiliation, and if available, zygosity was also included. The subjects (506 males, 606 females) had a mean age of 28.8 ± 3.7 years (range: 22 – 37 years).

### Identification of “bridged” central sulci

Image preprocessing, surface reconstruction and extractions of ‘Cortical Folds Graphs’ (CFG) representing the attributed relational graph structures of cortical sulci were performed using BrainVISA 4.5.0 and the Morphologist 2015 pipeline (Geffroy et al. 2011, Rivière et al. 2011). Preprocessing included transformation into AC-PC space and inhomogeneity correction. Cortical surfaces and CFGs were reconstructed, yielding 2224 hemispheric reconstructions. For each hemisphere, the CFGs of the CS were identified and independently examined by at least two out of three raters (J.S., a second rater, and A.M.M.), who worked independently on the dataset during three separate time periods. Criteria for identifying a ‘bridged CS’ included: (i) accurate identification of the CS in the T1-weighted images and grey matter surface reconstruction, (ii) correct localization of the ‘bridge’ in the lower boundary of the upper third of the CS associated with the hand knob structure at the precentral gyrus (Yousry et al. 1998), and (iii) completeness of the ‘bridged’ CS reaching the brain surface in the surface reconstruction with an incomplete CS depth profile either representing only the upper or the lower part of the discontinued CS. Each case was further reviewed and confirmed by R.S. and A.M.M. In one atypical case involving an unusual CS structure, functional MRI data of an HCP-YA finger tapping task was utilized to infer the precentral gyrus trajectory toward the interhemispheric cleft. Surface reconstructions for visualization were obtained using BrainVoyager Version 2.8.4. (Brain Innovation, Maastricht, The Netherlands).

### Determination of PPfpm heights and descriptive statistics

CFGs were obtained for 2165 left and right hemispheres with a continuous CS (Mangin et al. 2004, Regis et al. 2005). All CFG of the CS underwent visual quality control to ensure continuity, joining segmented units of the CS and excluding minor side branches for a single, continuous CS. Using BrainVISA ‘Sulcus Parametrization (2015), each CS CFG was parameterized to obtain a depth profile, which represents the distance between the brain surface boundary and the grey matter surface in the fundus of the CS at 101 equidistant positions along the CS mesh (Cykowski et al. 2008). Depth values were inverted assigning increasing negative values to increased depth, with the brain surface set to 0mm.

The absolute PPfpm height was calculated as the difference between the maximal depth value at positions > 45 and the next adjacent local maximum at position ≤ 45 (Cykowski et al. 2008, Müllen & Schweizer 2022). For each of the 1983 PPfpms, relative height was computed as the ration of the absolute PPfpm height to the respective maximum depth at positions > 45 of the individual CS. The median, mean (µ) and standard deviation (SD) of the relative PPfpm height distribution was then determined, with binning at 0.1 fractions of CS depth (fCSdepth).

To characterize the height distribution of the PPfpm in the HCP-YA dataset and assess the presence of extreme values, we modelled the distribution using truncated normal and beta distributions as plausible candidates for the distribution at hand, both of which accommodate values within the interval [0, 1]. First, we estimated the parameters of each distribution via maximum likelihood estimation (truncated normal distribution: mean, standard deviation, truncation thresholds set to 0 and 1; beta distribution: mean and a precision parameter). We then applied a non-parametric bootstrap (N=1,000 bootstraps) to sample with replacement from the original distribution, determining confidence intervals for the density distribution. For each bootstrap iteration, we sampled N values (where N = original sample size) from the original data with replacement, estimated distribution parameters, calculated the probability density function within the interval [0, 1] and established 95% confidence limits for the density distribution.

The analysis was implemented in R (version 4.0.3; R Core Team 2020). Maximum likelihood estimation was conducted using the function optim with method=“BFGS”, iterating with the parameters of the previous call until the log-likelihood difference between successive calls was less than 0.0001. For estimating the probability of the individual observations and determining the probability density distribution we used the functions dtruncnorm (truncated normal distribution) of the package truncnorm (version 1.0-8, Mersmann et al. 2018) or dbeta (beta distribution, stats package of R), respectively.

### Comparison of Heschl ‘s and the HCP-YA height distribution

Heschl categorized relative ‘deep winding’ heights, expressed as fraction of CS depth (fCSdepth) into three bins: (1) cases with a bridged CS, (2) heights between 1/3 - 5/6 fCSdepth and (3) height of 1/6 - 1/3 fCSdepth. He also noted that cases with lesser heights were not recorded. To facilitate comparison, we added a fourth category: cases with heights between 0 and 1/6 fCSdepth. The number of cases in this fourth category was calculated as the remainder of the total cases after accounting for those in the three reported categories.

To align the HCP-YA distribution, here reported in 100 bins, with Heschl’s distribution of only four bins, we summed the number of HCP-YA cases within Heschl’s bins. Given the considerable difference in relative ‘deep windings’ / PPfpm heights between the two distributions, we also adopted a second approach based on normal distribution parameters to provide a broader comparative assessment.

For the latter approach we applied descriptive statistics to Heschl’s bins. Since the first bin contains 93% of the cases, its range can be approximated by the mean (µ) and the standard deviation (σ) as −2σ < µ < +2σ or 4σ, representing the central portion of the normal distribution encompassing 95% of cases. As the range of the second bin equals the first, its upper boundary can be estimated as its lower boundary plus the width of 4σ: thus starting at µ + 2σ, ending at µ + 6α. The third category, three times wider than the first two with a range of 12σ and has a lower boundary at µ + 6σ with an upper boundary set at µ + 18σ. To apply these intervals to the HCP-YA data, we inserted the calculated mean (µ = 0.184 fCSdepth) and standard deviation (σ = 0.112 fCSdepth) as follows: the upper boundary of the first bin was µ + 2σ = 0.184 + 2 x 0.112 = 0.408 fCSdepth, of the second bin µ + 6σ = 0.184 + 6 x 0.112 = 0.856 fCSdepth and of the third bin µ + 18α = 2,2 fCSdepth, and with the last bin being irrelevant for the HCP-YA data (Tab.1).

**Table 1.** Boundaries of bins defined by Heschl (1877), expressed in statistical parameters of a normal distribution estimated from Heschl’s data [fCSdepth = fraction of CS depth]. Corresponding values for the HCP-YA data [fCSdepth = fraction of maximal CS depth] were calculated on the mean (µ) and standard deviation (0) obtained from the HCP-YA dataset.

| *Distribution* | *mean*<br>$\mu$ | *SD*<br>$\sigma$ | *boundary at*<br>$\mu + 2\sigma$ | *boundary at*<br>$\mu + 6\sigma$ | *boundary at*<br>$\mu + 18\sigma$ |
| --- | --- | --- | --- | --- | --- |
| Heschl data | -- | -- | 1/6 = 0.17 [fCSdepth] | 1/3 = 0.33 [fCSdepth] | 5/6 = 0.83 [fCSdepth] |
| HCP-YA data | 0.18 | 0.11 | 0.41 [fCSdepth] | 0.86 [fCSdepth] | 2.2 [fCSdepth] |

## Results

To replicate the historical prevalence of the ‘bridged’ CS in a large cohort (Heschl 1877), we conducted a visual inspection of 1112 MRI-based brain surface reconstructions from the HCP-YA dataset and present the detected cases. The heights of the deep annectant gyrus within the CS were obtained in 1983 hemispheres based on CS depth profiles and the resulting PPfpm height distribution compared with the ‘deep winding’ height distribution reported by Heschl. The high number of twins in the HCP-YA dataset allowed us to additionally assess the prevalence of the ‘bridged’ CS in both dizygotic and monozygotic twins, as well as a potentially higher prevalence within twin pairs.

### Prevalence of the “bridged” CS in the HCP-YA data set

Visual inspection of 1112 T1-weighted MRI brain surface reconstructions revealed nine cases with a unilateral ‘bridged’ CS yielding a prevalence of 0,8% (Fig.1A, Tab.1). The ‘bridged’ CS were in six cases located in the right hemisphere and in three cases in the left; six cases occurred in females (five in the right hemisphere, one in the left) and three in males (one in the right hemisphere, two in the left).

**Fig. 1.**
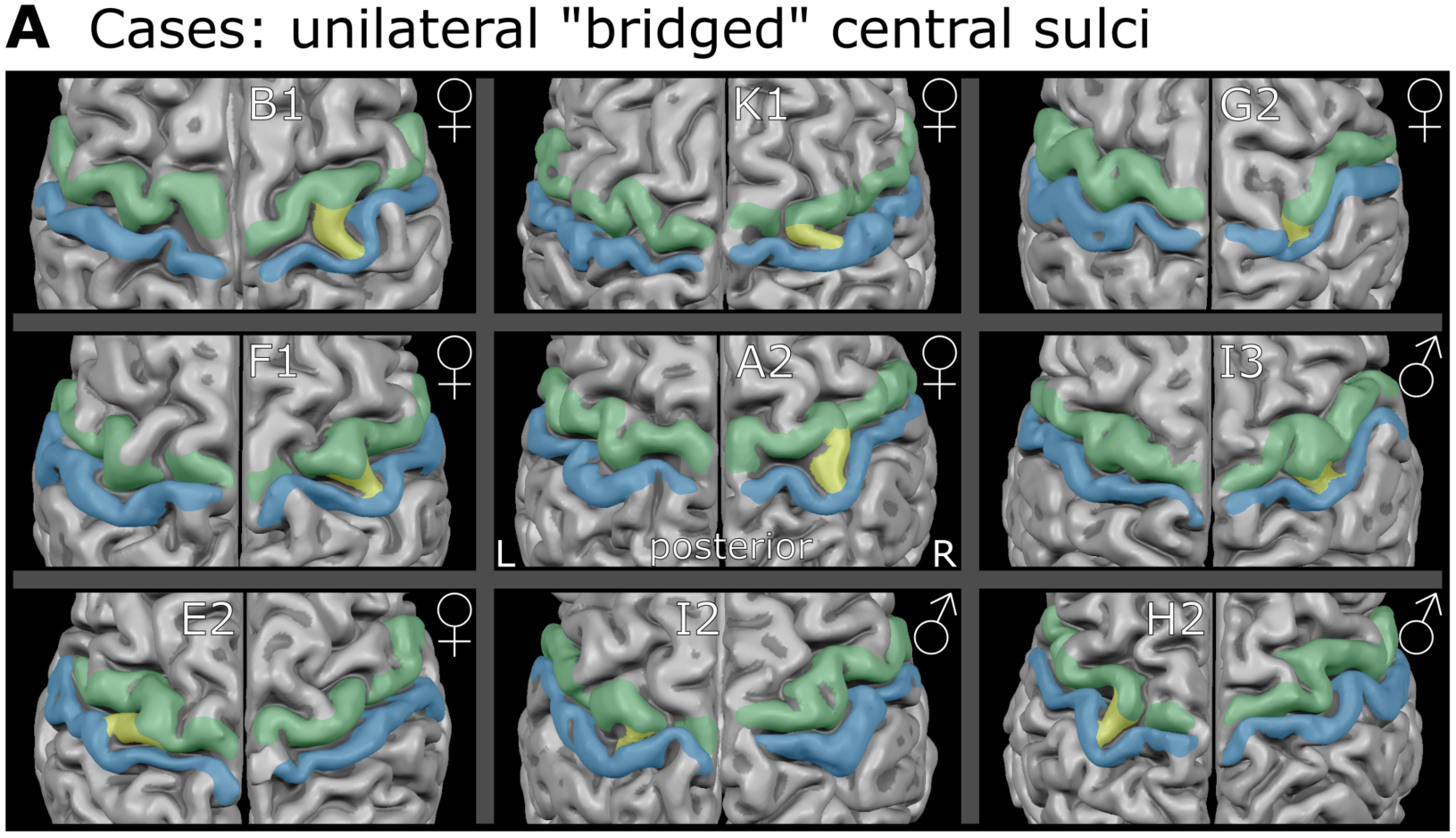
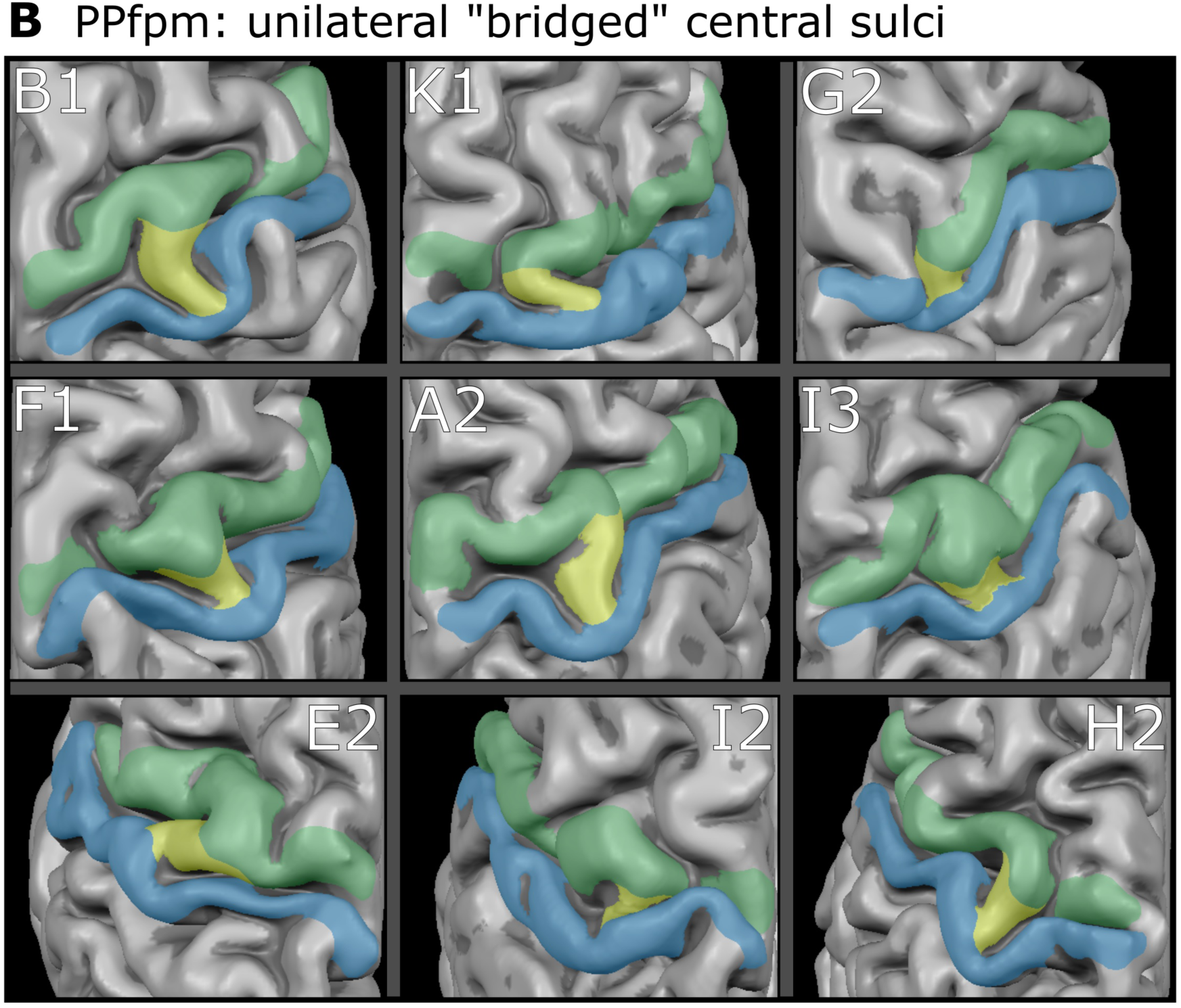
Unilateral “bridged” central sulci ***a*** Surface reconstructions of the central sulcus area for both hemispheres. Green = precentral gyrus; blue = postcentral gyrus; yellow = “bridged” central sulcus / PPfpm at the brain surface; ♀= female, ♂ = male; L = left, R = right; letters/numbers = family affiliation/position in sibling order **b** Enlargements of the unilateral ‘bridged’ central sulci shown in **a**

Comparison with Heschl’s (1877) findings indicates similarities in absolute counts, prevalence and hemispheric ratio, but a notable deviation in the gender distribution (Tab.1). Both datasets show comparable prevalence rates (HCP-YA = 0.8%; Heschl = 0.6%) and a consistent left-to-right-hemisphere ratio of 1:2. This underscores the similarities despite differences in the temporal and demographic composition of the datasets.

However, a marked difference emerges in gender distribution between the datasets (Table 2). In the HCP-YA dataset, the male-to-female ratio for ‘bridged’ CS cases is 1:2 (three males, six females), contrasting sharply with Heschl’s finding of 5:1 (five males, one female). This disparity is further reflected in the gender-specific prevalence: HCP-YA: males = 0.6%, females = 1% versus Heschl males = 0,8 %, females = 0.2%, accounting for differing male and female dataset sizes in each dataset.

**Table 2.** Number of ‘bridged’ central sulci and their distribution across hemispheres and gender in the HCP-YA dataset and the dataset of Heschl (1877)

| Parameters | HCP-YA dataset | Heschl (1877) dataset |
| --- | --- | --- |
| number of subjects in datasets | 1112<br>506 males / 606 females | 1087<br>632 males / 455 females |
| absolute number of ‘bridged’ CS | 9 | 6 |
| prevalence of ‘bridged’ CS | 0.8 % | 0.6% |
| hemispheres with ‘bridged’ CS | 3 left / 6 right | 2 left / 4 right |
| gender specific absolute number of ‘bridged’ CS | 3 males / 6 females | 5 males / 1 female |
| gender specific prevalence of ‘bridged’ CS | male: 0.6% / female: 1% | male: 0.8% / female: 0.2% |

### Individual Appearance of the Surface based PPfpm

Each of the nine cases of a “bridged” CS has a characteristic shape as well as a specific angle in crossing the CS (Fig.1b). In six cases, the PPfpm passes the CS perpendicular to its main trajectory, resulting in the PPfpm presenting as a direct and short connection (Fig.1b: B1, G2, F1, A2, I3, H2). In the remaining three cases, the PPfpm crosses diagonally (Fig.1b: K1, E2, I2) reaching from the more lateral postcentral gyrus towards medial where it connects to the precentral gyrus thus resulting in an elongated PPfpm with a shallow angle. The individual manifestations of these PPfpm seen on the brain surface might reflect the variability of the ‘deep winding’ PPfpm in the depth of the CS.

### PPfpm Height Distribution in the HCP-YA dataset

The relative heights of the PPfpm of the HCP-YA dataset result in a distribution with a median PPfpm height at approximately 3.97mm (0.167 fraction of maximal CS depth (fCSdepth)) and a mean PPfpm height of approximately 4.41mm ± 2.74mm (0.18 ± 0.11 SD fCSdepth) (Fig.2). The absolute heights given in millimeter are based on the original data obtained from the depth profiles. The higher mean value compared to the median reflects a skew in the distribution towards higher PPfpm heights, the most extreme case exhibiting a PPfpm height of approximately 17.5 mm (0.76 fCSdepth). The distribution itself can be better described by a beta distribution (log-likelihood: −1672.721) than by a truncated normal distribution (log-likelihood: −1682.148), although both theoretical distributions have a good match at the larger height values and a limited fit at small values up to 0.25.

**Fig. 2.**
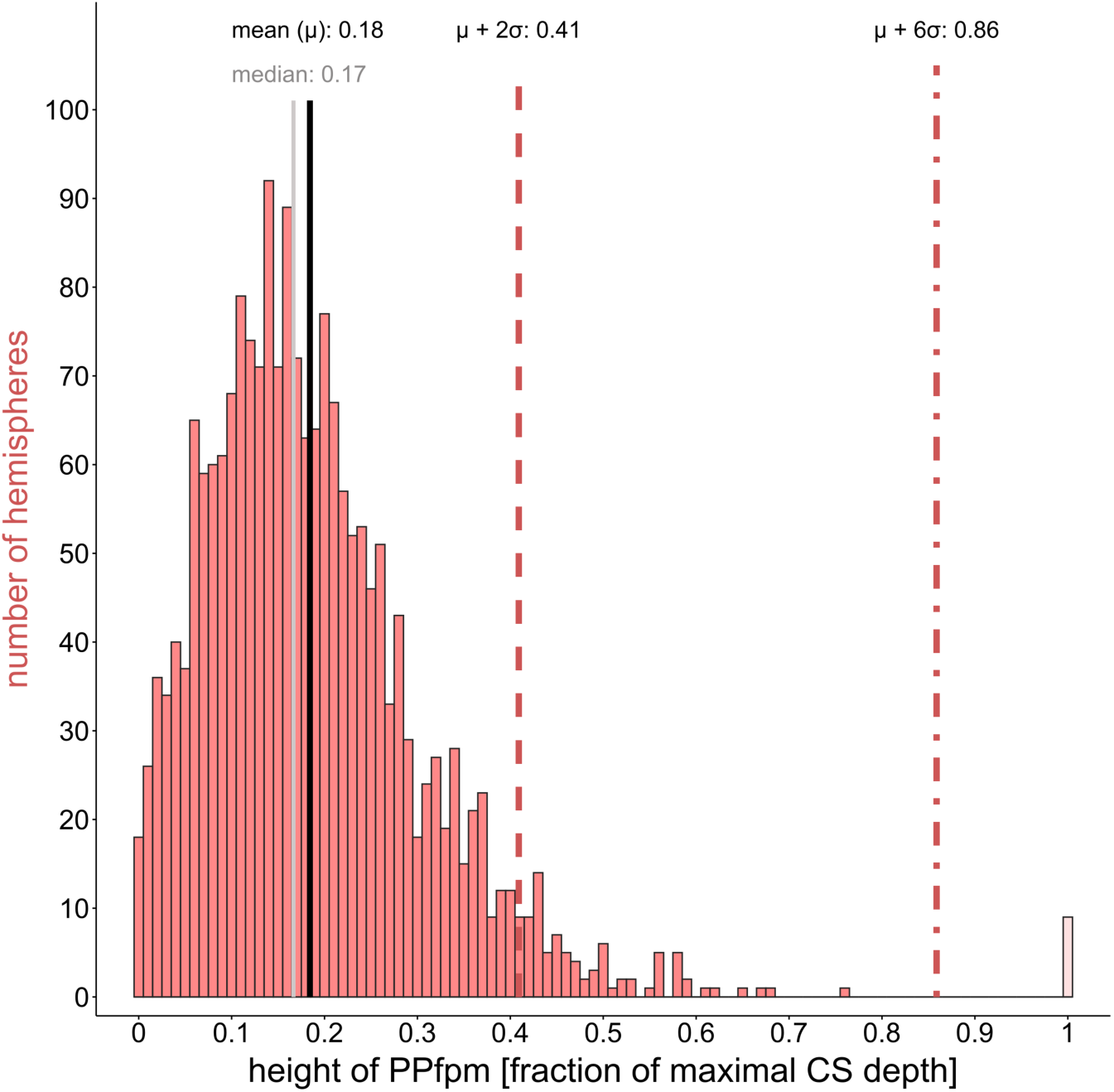
Height distribution HCP-YA data set: relative height of PPfpms (N =1983) expressed as fraction of maximal central sulcus depth (fCSdepth); 1.0 height = “bridged” central sulcus; black line = mean (µ); grey line = median; red line = standard deviation (σ), short-dashed line = µ+2σ; dash-dotted line = µ+6σ.

### ‘Deep Winding’ Height Distribution in the Heschl dataset

Heschl’s measurements indicate that the majority of the ‘deep winding’ are of low height, but also reveal a positively skewed distribution, with ‘deep winding’ heights to just below the brain surface. The relative heights of the ‘deep winding’ are reported in four bins of varying widths (Fig.3). The largest proportion, 93%, falls below 1/6 fCSdepth, labeled by Heschl as ‘undeterminable minor height’. An additional 4% of the cases are within the “1/6 – 1/3” fCSdepth range, while the remaining 3% occupy the “1/3 – 5/6” fCSdepth range, covering ‘deep winding’ heights extending to just below the brain surface. The final bin comprises six cases of unilateral ‘bridged’ CS, were the ‘deep winding’ is present at brain surface level (Fig.3, Table 3)

**Fig. 3.**
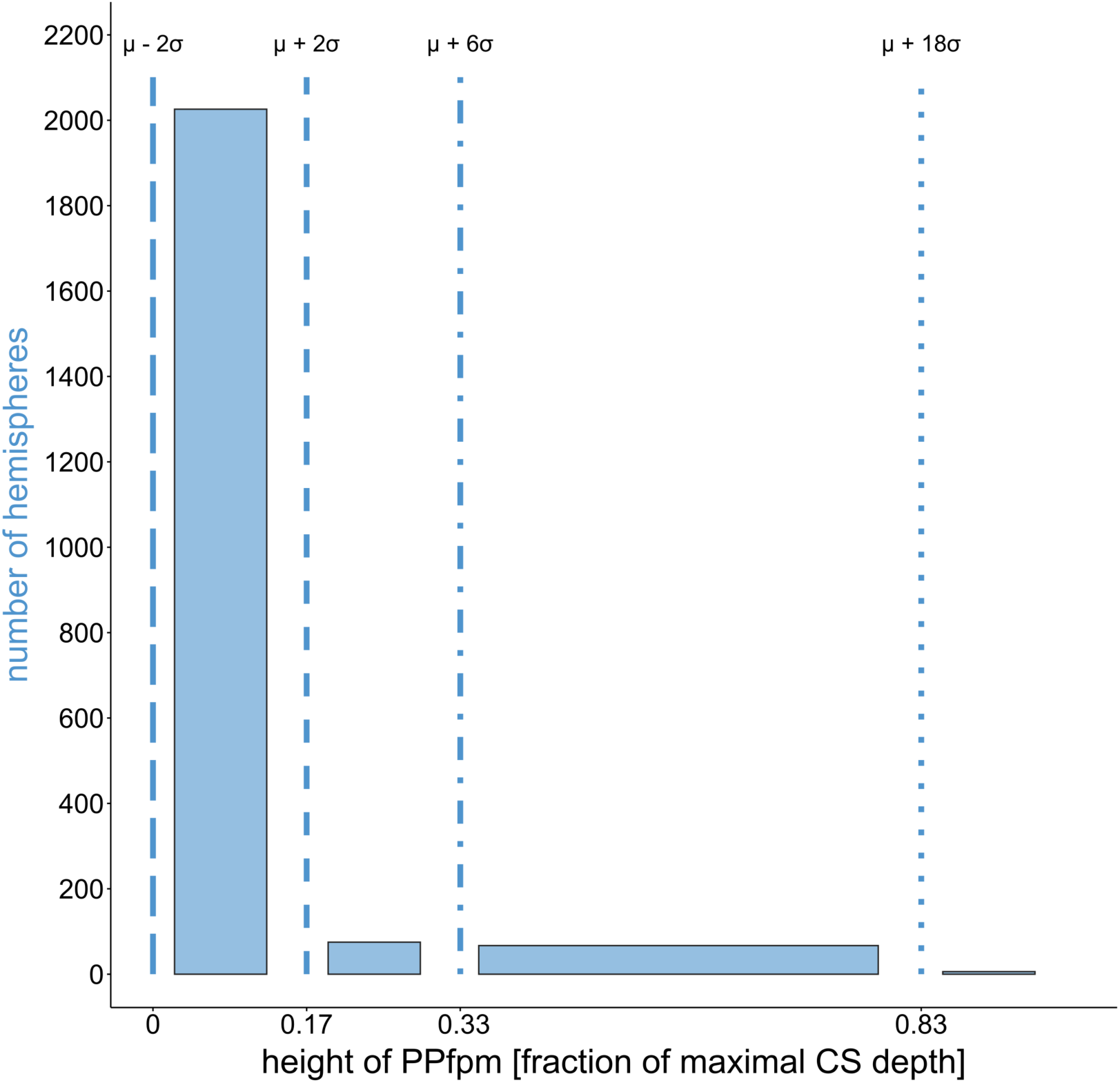
Height distribution Heschl (1877) data set: relative height of ‘deep winding’ (N = 2174) as a fraction of maximal central sulcus depth (fCSdepth); blue lines = boundaries of the bins defined by Heschl (1877) at 1/6 (0.17), 1/3 (0.33) and 5/6 (0.83) fCSdepth.

**Table 3.** Comparison of ’deep winding’ / PPfpm height distributions between Heschl (1877) and HCP-YA dataset.

|  |  | *Heschl 0 - 1/6*<br>fraction of CS depth |  | *Heschl 1/6 – 1/3*<br>fraction of CS depth |  | *Heschl 1/3 – 5/6*<br>fraction of CS depth |  |
| --- | --- | --- | --- | --- | --- | --- | --- |
|  |  | *HCP-YA 0 – 0.17*<br>fraction of CS depth |  | *HCP-YA 0.17 – 0.33*<br>fraction of CS depth |  | *HCP-YA 0.33 – 0.83*<br>fraction of CS depth |  |
| *dataset* | *Total number of PPfpm* | *total PPfpm* | *percent PPfpm* | *total PPfpm* | *percent PPfpm* | *total PPfpm* | *percent PPfpm* |
| Heschl | 2168 | 2026 | 93% | 75 | 4% | 67 | 3% |
| HCP-YA | 1983 | 992 | 50% | 780 | 39% | 211 | 11% |

### Comparison of the Heschl’s ‘Deep Winding’ and HCP-YA PPfpm Distributions

The comparison of the HCP-YA PPfpm height distributions, resampled in the bins of Heschl, with Heschl’s ‘deep winding’ height distribution, reveal generally greater PPfpm heights and a broader distribution range in the HCP-YA dataset (Fig.4).

**Fig. 4.**
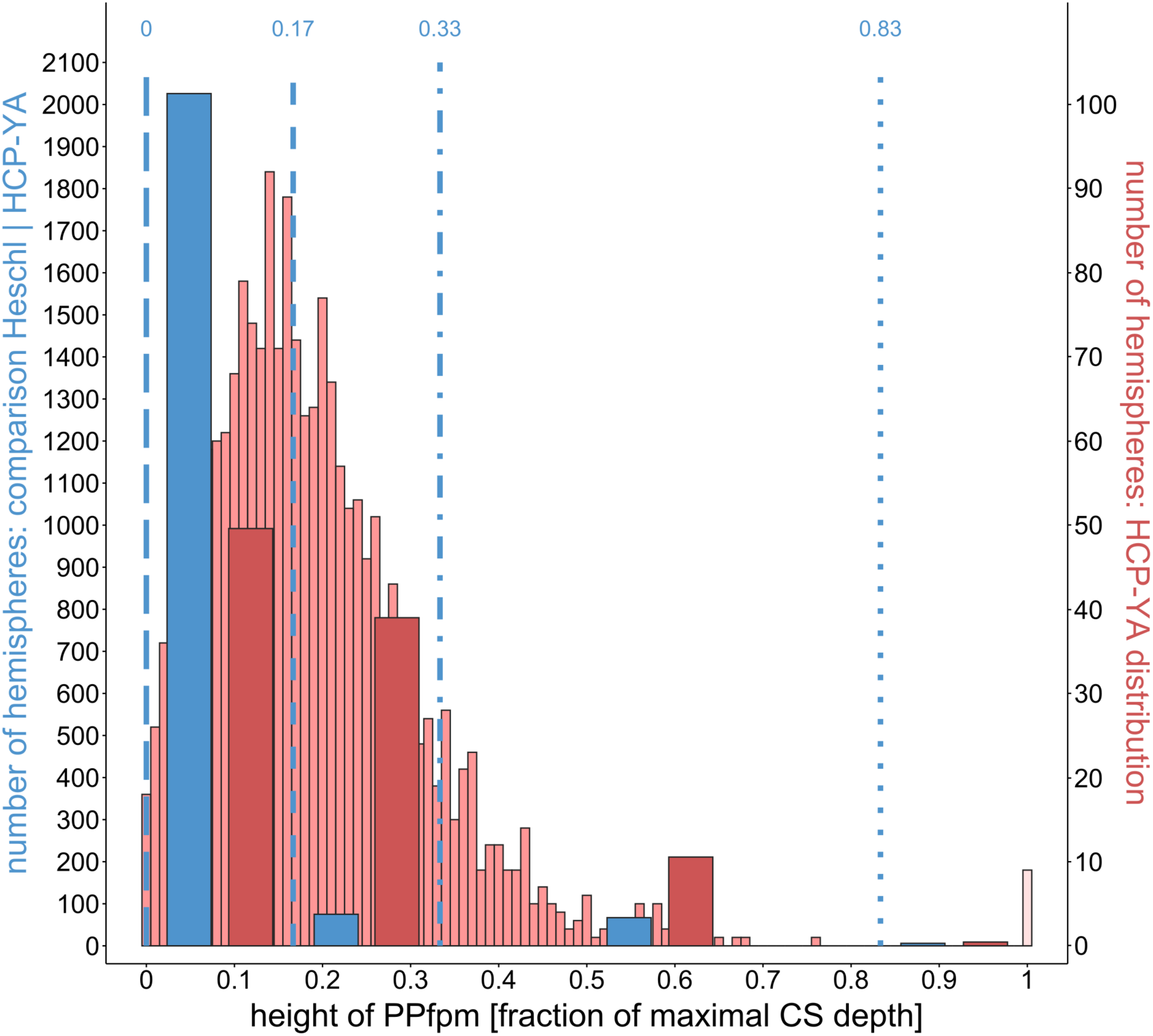
Comparison of HCP-YA and Heschl (1877) distribution based on relative PPfpm height (fraction of maximal central sulcus depth, fCSdepth) using Heschl’s binning. The left Y-axis represents counts from Heschl’s data set and HCP-YA data within Heschl’s bins, the right Y-axis shows HCP-YA data counts in the binning applied to the HPC-YA data set. Blue lines = Heschl’s ‘deep winding’ heights as fCSdepth: long-dashed line = 0 fCSdepth; short-dashed line = 0.17 (1/6) fCSdepth; dash-dotted line = 0.33 (1/3) fCSdepth; dotted line = 0.83 (5/6) fCSdepth

The HCP-YA data show 50% of PPfpms in the first bin, 39% in the second and 11% in the third, widest bin (Table 3). This contrasts with Heschl’s data, where 93% of the ‘deep windings’ falls within first bin. The HCP-YA distribution thus displays nearly twice the spread, with 89% of PPfpm heights dispersed across the first two bins. This broader distribution with generally larger PPfpm heights in the HCP-YA dataset likely results from differences in methods used to obtain PPfpm heights.

This noticeable shift in the HCP-YA distribution toward higher PPfpm height values poses limitations for directly comparing the two datasets. To address this, we translated the boundaries of Heschl’s bins from ‘fractions of CS depth’ into statistical terms, using mean ± standard deviation approximations based on the normal distribution. We then converted the HCP-YA data into corresponding bins based on this approximation.

This approach highlights the considerable similarities between the two distributions. For the first bin, similarity is expected due to the properties of the normal distribution. In the second bin, both distributions exhibit a comparable skew toward higher PPfpm heights, with deviations in heights reaching to up to six standard deviations. The third bin, due to narrower width in Heschl’s distribution and a smaller estimated standard deviation, includes only the highest ‘deep windings’. In both datasets, the ‘bridged’ CS cases are classified in the fourth bin (Fig.5, Table 4).

**Fig. 5.**
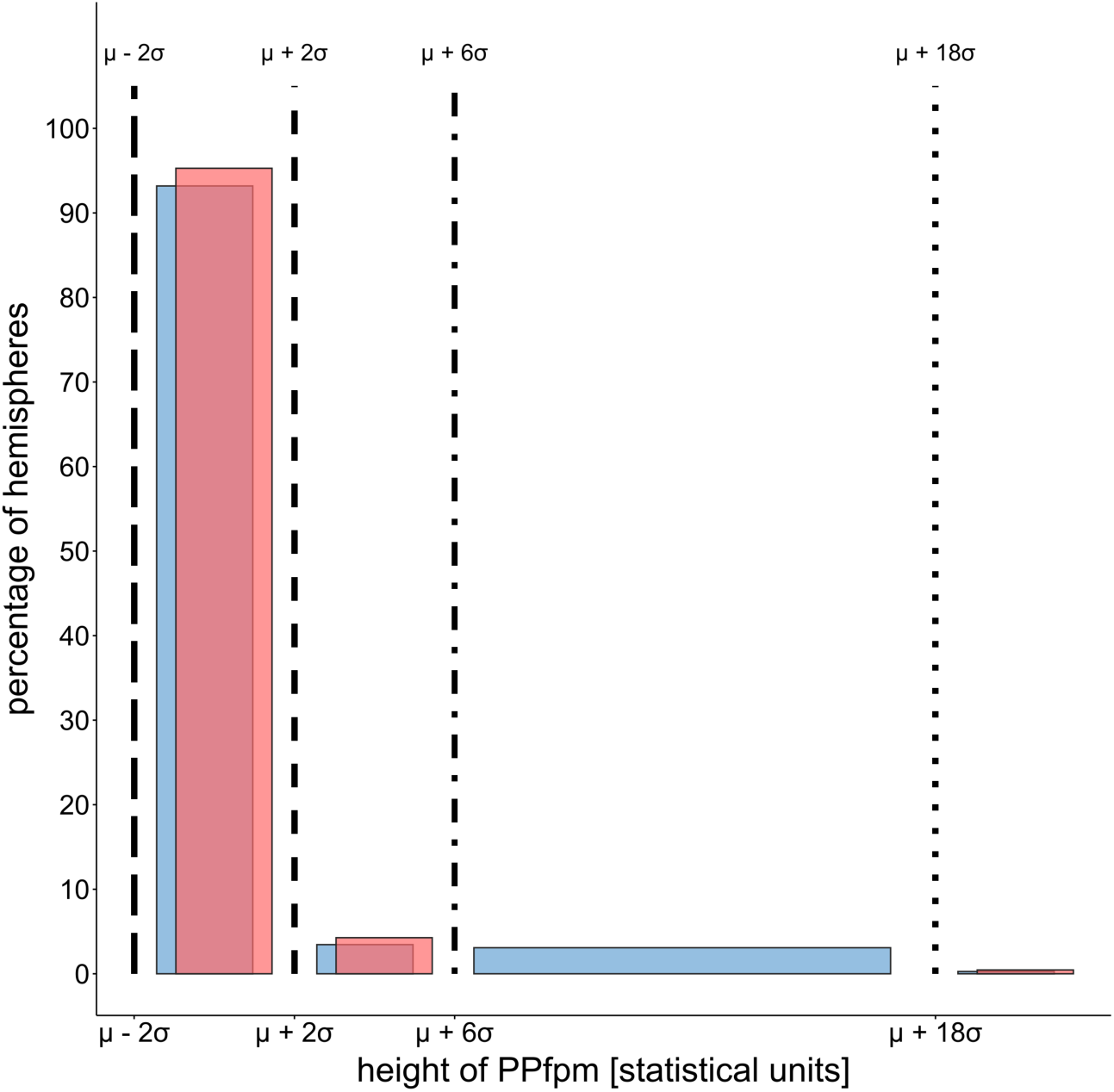
Comparison of HCP-YA and Heschl (1877) ‘deep winding’ height distributions based on normal distribution parameters estimated from Heschl’s bins. µ = acquired (HCP-YA) or estimated (Heschl) mean, σ = acquired (HCP-YA) or estimated (Heschl) standard deviation. Black lines = Heschl’s bins transformed into statistical parameters: long-dashed line = µ-2σ; short-dashed line = µ+2σ; dash-dotted line = µ+6σ; dotted line = µ+18σ

**Table 4.** Comparison of case frequencies in Heschl (1877) and HCP-YA ‘deep winding’ / PPfpm height distributions, using statistical parameters to define Heschl’s bins.

| Distribution | Heschl | HCP-YA |
| --- | --- | --- |
| 1. bin [% cases] | 93.2% | 95.7% |
| 2. bin [% cases] | 3.5% | 4.3% |
| 3. bin [% cases] | 3.1% | -- |
| 4. bin [# of 'bridged' CS] | 6 | 9 |

The observed skew toward larger ‘deep winding’ and PPfpm heights in both distributions supports Heschl’s hypothesis: “If this depth winding were a typical one, the imaginary bridging would have to be located at the point where it is usually found in a less developed form. … It must now be further concluded that (..the deep winding..) must occur in all degrees of formation and that a case of total interruption of the central sulcus by the same will not be long in coming with systematic searches.”

### Prevalence and Occurrence of a ‘bridged’ Central Sulcus in Non-Twin and Twin Siblings

The dataset suggests a higher prevalence of a ‘bridged’ CS in individuals who are dizygotic or monozygotic twins, though not specifically within twin pairs. The nine cases with a ‘bridged’ CS are evenly distributed across the three sibling groups: three cases among non-twins, three among monozygotic twins, and three among dizygotic twins (Table 5). Given the composition of 59.1% non-twins, 15.3% dizygotic twins, 25.6% monozygotic twins, this results in a ‘bridged’ CS prevalence of 0.5% for non-twins, 1.8% for dizygotic twins, and 1.1% for monozygotic twins. These figures may suggest a higher likelihood of a ‘bridged’ CS among twin siblings.

**Table 5.** Prevalence of a ‘bridged’ CS in twin and non-twin siblings.

|  | Non-twins | Dizygotic Twins | Monozygotic Twins |
| --- | --- | --- | --- |
| Total number HCP-YA | 657 | 170 | 285 |
| Percentage HCP-YA | 59.1% | 15.3% | 25.6% |
| Incidence ‘bridged’ CS | 3 | 3 | 3 |
| Prevalence ‘bridged’ CS | 0.5% | 1.8% | 1.1% |

To explore the potentially higher familial prevalence of ‘bridged’ CS cases, the nine instances were analyzed in relation to non-twin and twin siblings who also participated in the HCP-YA study. Among the three non-twin cases, none of their twin siblings exhibited a ‘bridged’ CS. This pattern is consistent among the three dizygotic twins, and one monozygotic twin with each a ‘bridged’ CS, whose siblings also did not exhibit a ‘bridged’ CS (Fig.6).

**Fig. 6.**
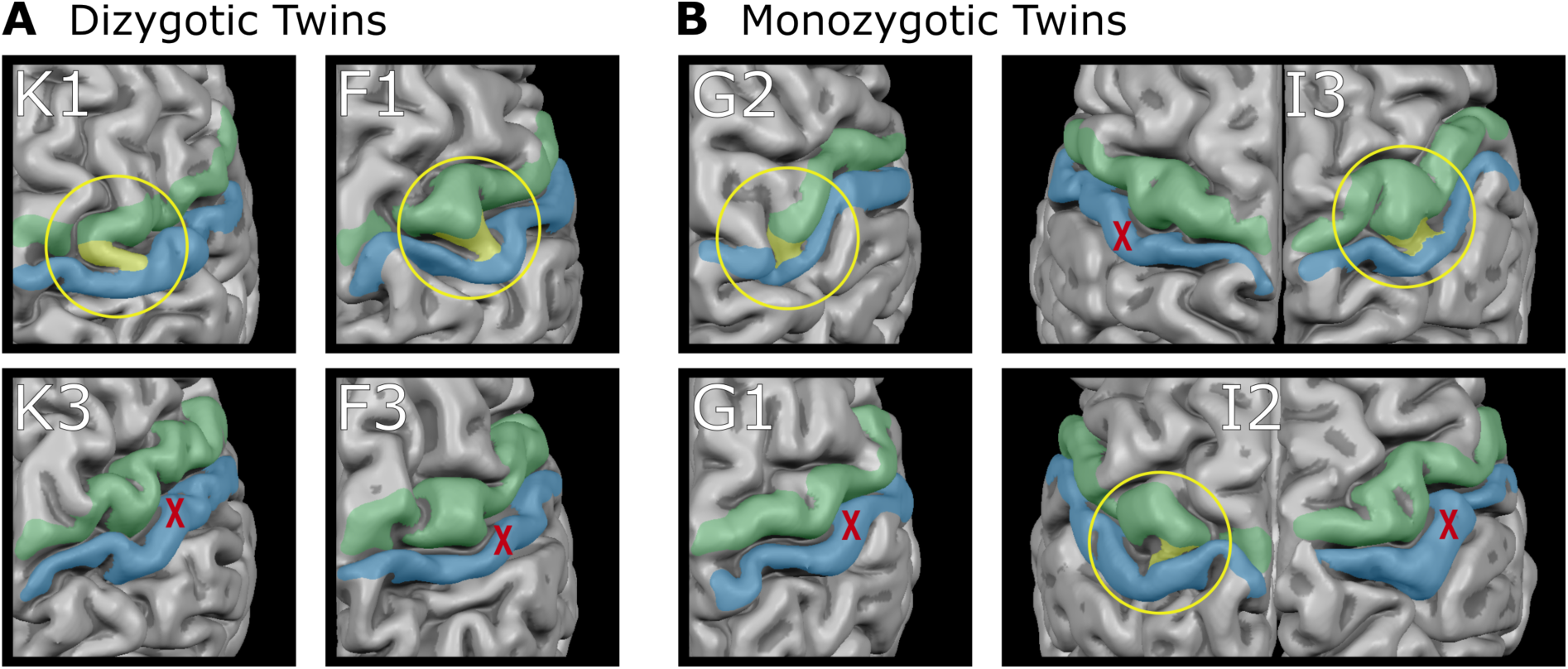
Unilaterale “bridged” central sulci in twin pair siblings. Surface reconstructions of the central sulcus area are shown for **a** Dizygotic Twins **b** Monozygotic Twins Green = precentral gyrus; blue = postcentral gyrus; yellow = “bridged” central sulcus - PPfpm at the brain surface; letters/numbers = family affiliation, yellow circle: “bridged” central sulcus - PPfpm at the brain surface, red cross: absence of “bridged” central sulcus - PPfpm at the brain surface in the same hemisphere of the twin sibling

The only exception of what involves the remaining two cases of a ‘bridged’ CS in a monozygotic twin pair: While both twins display a unilateral ‘bridged’ CS, they occur in opposite hemispheres. Given the independence of hemispheric PPfpm height, this likely indicates an unrelated, coincidental, occurrence within this twin pair. This interpretation is further supported by the overall layout of the central sulci in these monozygotic twins, which show the same variability and dissimilarity as observed in other monozygotic or dizygotic twin sibling (Fig.6) and in the brains of the other seven cases with a ‘bridged’ CS (Fig.1).

## Discussion

We replicated Heschl’s approach to determine the prevalence of the ‘bridged’ CS and the height distribution of the ‘deep winding’ using a contemporary dataset of comparable size. Our visual search of the MRI-based surface reconstructions yielded a similar prevalence of unilateral ‘bridged’ CSs, suggesting a consistent prevalence, regardless of dataset composition. The MRI-based assessment of the HCP-YA PPfpm heights refines Heschl’s original findings and reveal generally higher height values and a broader distribution. Nevertheless, Heschl’s finding of a pronounced skew of the ‘deep windings’ towards larger heights was confirmed in the HCP-YA dataset, further supporting the relationship between the ‘deep winding’ / PPfpm and the ‘bridged’ CS. While monozygotic and dizygotic twins exhibited a slightly higher prevalence of a ‘bridged’ CS, no cases of congruous unilateral ‘bridged’ CS were found in monozygotic twin pairs.

### Prevalence of ‘bridged’ CS

The decision to use a visual search on the MRI-based whole brain surface reconstructions, assumed to parallel the historical approach on brain specimens, provides a methodologically unbiased perspective of the results. The similar prevalence rates of the two datasets - healthy young adult volunteers in the HCP-YA dataset and the deceased patients from the general hospital in Vienna in 1876 – strongly suggest that factors such as age, health, disease or lifestyle do not substantially influence the occurrence of a ‘bridged’ CS across samples of historical European and contemporary North American origin. While the slight difference in prevalence, 0.6% in the historical dataset versus 0.9% in the contemporary dataset, may be considered noteworthy, it could be attributed the higher proportion of twins in the HCP-YA dataset, as twins exhibit a higher prevalence of ‘bridged’ CA compared to non-twins (1.45% vs 0.5%). However, large-scale studies of both twin and non-twin populations are needed to confirm this finding. The same applies to the comparable prevalence of unilateral ‘bridged’ CS in the left or right hemisphere, as well as the dissimilarity of prevalence between males and females in the two datasets. Although the large dataset size provides a robust basis for identifying ‘bridged’ CS cases, the relatively small number of detected cases is insufficient to statistically determine significant difference in prevalence by sex or hemisphere.

The methodological similarity of the visual search is limited by the difference in the observational techniques. Heschl likely examined whole brain specimens directly, whereas modern analysis relies on viewing MRI-based 3D brain surface reconstructions on a monitor. One advantage of T1-weighted structural MRI is the ability to visualize the entire brain using sectional planes from any direction and angle. However, the method also has drawbacks, such as latent imaging artefacts, which can introduce imprecisions in brain surface reconstruction. These inaccuracies may affect the determination of the ‘completeness’ of a ‘bridged’ CS, particularly whether it is present at the brain surface or rather a unusually high “deep winding” PPfpm. We followed Heschl’s criteria, defining a deep winding being greater than 5/6 of the CS depth for characterizing a ‘bridged’ CS. However, the MRI surface reconstructions of the selected cases still show the remaining degree of variability.

A second limitation relates to the ambiguity in the number of cases identified in the HCP-YA dataset, likely influence by the general and specific expertise of the raters as well as cases of highly atypical CS anatomy. Heschl, renowned for his work on the anatomy of the auditory cortex, possessed extensive anatomical knowledge and experience, which informed his judgements. Given that this level of expertise is now difficult to attain, each CS was visually inspected by at least two of three independent raters. The value of this approach is evident in the increasing number of ‘bridged’ CS cases reported in successive conference abstracts, reflecting not only a growing number of analyzed brains but also the raters’ enhanced anatomical expertise (Schweizer et al. 2018, Schweizer et al. 2018, Müllen & Schweizer 2022). However, some ambiguity persists. Mangin and colleagues (2019), using an algorithm to detect the ‘bridged’ CS in the HCP-YA dataset, reported the same number of cases as we did previously (Schweizer et al. 2019), but with three differing individual cases of a ‘bridged’ CS. This discrepancy can largely be attributed to extremely deviant CS anatomies, which are naturally present in large datasets and may complicate the clear identification of a ‘bridged’ CS. Such cases could benefit from incorporating additional and more detailed anatomical criteria. Nevertheless, these variations in individual cases should not undermine the current approaches, as establishing a definite prevalence of the ‘bridged’ CS would require precise anatomical criteria, which goes beyond the scope of this replication.

### PPfpm Height Distribution

Contrasting the visual search to detect cases of ‘bridged’ CS, the heights of the PPfpm were estimated by the automized extraction of the MRI-based CS depth profile (Cykowski et al. 2008) and a consequential automized determination of two landmarks related to the PPfpm (Müllen & Schweizer 2022). We interpret the observed shift toward larger heights and the increased range of the PPfpm height distribution as a refinement resulting from contemporary methods rather than dataset differences.

A limitation of the automated PPfpm height acquisition lies in the precision of estimation, reflecting the challenge of translating complex individual anatomy into objective, comparable metrics. Each algorithm captures specific anatomical aspects based on certain assumptions, introducing potential errors. But a certain degree of errors may also have occurred in Heschl’s measurements. As the aim of this replication was not to measure absolute PPfpm heights but to compare the shape of the distributions, these measurement errors are considered minor.

Despite differences in ‘deep winding’ PPfpm height and distribution width, the similarity between the HCP-YA PPfpm and the historical ‘deep winding’ height distributions, both showing a skew toward higher height values, support the validity of our findings. The increased distribution width becomes particularly apparent when resampling of the HCP-YA data to match Heschl’s height categories. To adjust for this, we applied normal distribution principles to convert historical bin boundaries into multiples of approximated standard deviations based on the historical ‘deep winding’ height frequency in Heschl’s first category. This unconventional approach arises from our belief that Heschl was not only an exceptional anatomist, but also possessed statistical insight. We respectfully assume that he may have been familiar with the concept of the normal distribution (Gauss 1809), and designed his categories with intent, viewing 93% of cases with “indeterminate heights” as typical “deep windings” and the remaining 7% as exceptionally high.

Converting both the historical and contemporary PPfpm height distributions into units of their respective standard deviations along with resampling adjusted to Heschl’s the category boundaries enabled a comparison independent of the differing distribution widths. This adaptation confirms not only the presence of extreme ‘deep winding’ PPfpm heights, but also Heschl’s proposition that the ‘bridged’ CS represents an extreme form of the ‘deep winding’ PPfpm.

### Twins

The high percentage of twins in the HCP-YA dataset allowed us to identify an elevated prevalence of ‘bridged CS’ in monozygotic and dizygotic twins, likely contributing to the slight increase in ‘bridged CS’ cases in the HCP-YA dataset compared to the historical dataset (HCP-YA: 5 twin individuals among the 9 cases; Heschl: 6 cases). However, this increased prevalence was not accompanied by a congruent unilateral ‘bridged’ CS for both individuals of monozygotic twin pairs. The dataset includes one monozygotic twin pair where both co-twins exhibit a ‘bridged’ CS, but in different hemispheres. Based on the general differences of CS in the two hemispheres, we interpret this occurrence as coincidental, facilitated by the elevated prevalence in twins. Both findings, the increased prevalence in monozygotic and dizygotic twin individuals and the missing co-occurrence in monozygotic twin pairs suggests that the occurrence of a ‘bridged’ CS is less influenced by genetic makeup and more affected by environmental factors.

Structural MRI studies of monozygotic twins show high correlation in general brain measures, e.g. volume of white matter, grey matter and corpus callosum, indicating genetic influence, whereas surface measures like gyral or sulcal patterns exhibit less similarity and greater environmental influence (Bonan et al. 1997, White et al. 2002). A study specifically examining the CS in adult monozygotic twins found a weak but significant correlation in homologous CS suggesting that CS morphology is only partially genetically determined and may be influenced by prenatal factors as early as the fifth month (Bonan et al. 1997). This aligns to the time of origin of the ‘bridged CS’ in the adult brain, being a remnant of a transitional stage around the 4^th^ month of the regular fetal CS development (Cunningham 1890, 1892), which typically recedes into the fundus of the CS as the common ‘deep winding’ PPfpm. The elevated prevalence in twins as well as the lack of co-occurrence in monozygotic twin pairs may therefore be attributed to the perinatal environmental of a twin pregnancy, possibly contributing to an early halt in the development of the CS, thus resulting in the ‘bridged’ CS.

Given that twins are known to face an increased risk of congenital development defects in brain morphogenesis (Park et al. 2020), it is important to emphasize that the incomplete development observed in the ‘bridged’ CS does not confer any apparent functional advantage or disadvantage. The increased grey matter area and white matter connections between the pre- and postcentral gyrus at the hand/finger area across the CS represent a structural variation that has no known impact on motor function or somatosensory perception.

## Conclusion

The present study successfully replicated Heschl’s historical survey of the rare ‘bridged’ CS and the heights of the ‘deep winding’ by examining MR-images from the similarly sized HCP-YA dataset. Visual inspection of brain surface reconstructions revealed a comparable prevalence of ‘bridged’ CS, with a slightly higher number of cases, likely reflecting the elevated prevalence observed in monozygotic and dizygotic twins in the HCP-YA dataset. The absence of concordance in monozygotic twin pairs suggests that twin pregnancy may act as an environmental factor in the formation of the ‘bridged’ CS. The height distribution of the PPfpm in the contemporary HCP-YA dataset shows a higher median and a broader range compared to Heschl’s findings. However, the distribution retained a similar skew toward larger PPfpm height values, as originally described for the ‘deep windings’, a pattern explicitly confirmed through data transformation. This successful replication corroborates Heschl’s hypothesis that the ‘deep winding’ and the ‘bridged’ CS represent different manifestations of the same anatomical structure. It also highlights Heschl’s remarkable anatomical expertise, combining meticulous observations with the innovative application of statistical approaches to investigate both, the rare and the common. By combining historical insights with contemporary methodologies, this study refines the anatomical understanding of the CS, facilitating more precise and comprehensive anatomical descriptions in future research.

## Author Contributions

RS: Conceptualization, Methodology, Validation, Resources, Writing - Original Draft, Writing - Review & Editing, Supervision, Project Administration, Funding Acquisition. AMM: Methodology, Software, Validation, Formal Analysis, Data Curation, Writing - Review & Editing, Visualization, Funding Acquisition. JS: Software, Formal Analysis, Reviewing

## Acknowledgements

We acknowledge funding by the Leibniz Association through an Outgoing Grant (LSC_OG2016_02) from the Leibniz ScienceCampus Primate Cognition (SAS-2015-DPZ-LWC) to R.S. and by the International Max Planck School for Neurosciences at the Georg-August University of Göttingen and by the German Academic Scholarship Foundation (Oct 2018 – March 2019) to A.M.M.

Data were provided by the Human Connectome Project, WU-Minn Consortium (Principal Investigators: David Van Essen and Kamil Ugurbil; 1U54MH091657) funded by the 16 NIH Institutes and Centers that support the NIH Blueprint for Neuroscience Research; and by the McDonnell Center for Systems Neuroscience at Washington University.

We extend our gratitude to Roger Mundry for his expert statistical analysis of the HCP-YA height distribution and Abdolmoein Esghaei for reminding us of the twins in the HCP-YA dataset and thereby inspiring the additional analysis.

## References

Alkadhi, H., & Kollias, S. S. (2004). Pli de passage fronto-parietal moyen of Broca separates the motor homunculus. American Journal of Neuroradiology, 25(5), 809–812.

Block, J., & Kuckertz, A. (2018). Seven principles of effective replication studies: strengthening the evidence base of management research. Management Review Quarterly, 68(4), 355–359. 10.1007/s11301-018-0149-3

Boling, W., Olivier, A., Bittar, R. G., & Reutens, D. (1999). Localization of hand motor activation in Broca’s pli de passage moyen. Journal of Neurosurgery, 91(6), 903–910. 10.3171/jns.1999.91.6.0903

Boling, W. W., & Olivier, A. (2004). Localization of hand sensory function to the pli de passage moyen of Broca. Journal of Neurosurgery, 101(2), 278–283. 10.3171/jns.2004.101.2.0278

Bonan, I., Argenti, A. M., Duyme, M., Hasboun, D., Dorion, A., Marsault, C., & Zouaoui, A. (2014). Magnetic Resonance Imaging of Cerebral Central Sulci: a Study of Monozygotic Twins. Acta geneticae medicae et gemellologiae: twin research, 47(2), 89–100. 10.1017/S000156600000026X

Brandt, M. J., Ijzerman, H., Dijksterhuis, A., Farach, F. J., Geller, J., Giner-Sorolla, R., Grange, J. A., Perugini, M., Spies, J. R., & van ’t Veer, A. (2014). The Replication Recipe: What makes for a convincing replication? Journal of Experimental Social Psychology, 50, 217–224. 10.1016/j.jesp.2013.10.005

Broca, P. (1888). Memoires d’Anthropologie. Reinwald.

Cunningham, D. J. (1890). The Fissure of Rolando. Journal of Anatomy and Physiology, 25, 1–23.

Cunningham, D. J. (1892). Contribution to the Surface Anatomy of the Cerebral Hemispheres. Academy House.

Cykowski, M. D., Coulon, O., Kochunov, P. V., Amunts, K., Lancaster, J. L., Laird, A. R., Glahn, D. C., & Fox, P. T. (2008). The Central Sulcus: an Observer-Independent Characterization of Sulcal Landmarks and Depth Asymmetry. Cereb. Cortex, 18(9), 1999–2009. 10.1093/cercor/bhm224

Eberstaller, O. (1884). Zur Oberflächenanatomie der Grosshirnhemisphären. Wien. Med. Blätter 7(16), 479–482.

Ecker, A. (1869). Die Hirnwindungen des Menschen. Friedrich Vieweg und Sohn.

Gauss, C. F. (1809). Theoria motus corporum coelestium in sectionibus conicis solem ambientium. F. Perthes et I.H. Besser.

Geffroy, D., Rivière, D., Denghien, I., Souedet, N., Laguitton, S., & Cointepas, Y. (2011). BrainVISA: a complete software platform for neuroimaging. Python in Neuroscience Paris.

Heschl, R. (1877). Tiefen-Windungen des menschlichen Grosshirns und die Überbrückung der Zentralfurche. Wiener Medizinische Wochenzeitschrift, 41, 987–988.

Mangin, J.-F., Le Guen, Y., Labra, N., Grigis, A., Frouin, V., Guevara, M., Fischer, C., Rivière, D., Hopkins, W. D., Régis, J., & Sun, Z. Y. (2019). “Plis de passage” Deserve a Role in Models of the Cortical Folding Process. Brain Topography, 32(6), 1035–1048. 10.1007/s10548-019-00734-8

Mangin, J. F., Riviere, D., Cachia, A., Duchesnay, E., Cointepas, Y., Papadopoulos-Orfanos, D., Collins, D. L., Evans, A. C., & Regis, J. (2004). Object-based morphometry of the cerebral cortex. Ieee Transactions on Medical Imaging, 23(8), 968–982. 10.1109/TMI.2004.831204

Mersmann, O., Trautmann, H., Steuer, D., & Bornkamp, B. (2018). truncnorm: Truncated Normal Distribution. R package version 1.0-8

Müllen, A.M. & Schweizer, R. (2022). Relating Depth Profiles of the Reconstructed Human Central Sulcus to Macroscopic Anatomical Landmarks. NEURIZONS2022, Goettingen, Germany.

Park, K. B., Chapman, T., Aldinger, K. A., Mirzaa, G. M., Zeiger, J., Beck, A., Glass, I. A., Hevner, R. F., Jansen, A. C., Marshall, D. A., Oegema, R., Parrini, E., Saneto, R. P., Curry, C. J., Hall, J. G., Guerrini, R., Leventer, R. J., & Dobyns, W. B. (2021). The spectrum of brain malformations and disruptions in twins [https://doi.org/10.1002/ajmg.a.61972]. American Journal of Medical Genetics Part A, 185(9), 2690–2718. 10.1002/ajmg.a.61972

Regis, J., Mangin, J.-F., Ochiai, T., Frouin, V., Riviere, D., Cachia, A., Tamura, M., & Samson, Y. (2005). “Sulcal Root” Generic Model: a Hypothesis to Overcome the Variability of the Human Cortex Folding Patterns. Neurologia Medico-Chirurgica, 45(1), 1–17. 10.2176/nmc.45.1

Retzius, G. (1896). Das Menschenhirn. Studien in der makroskopischen Morphologie. Kgl. Buchdr. P.A. Norstedt & Söner.

Retzius, G. (1901). Zur Frage von den sogenannten transitorischen Furchen des Menschenhirnes. Anat. Anz., 19, 91–92.

Rivière, D., Geffroy, D., Denghien, I., Souedet, N., & Cointepas, Y. (2011). Anatomist: a python framework for interactive 3D visualization of neuroimaging data. Python in Neuroscience workshop, Paris.

Schweizer, R., Graetz, H., Stropel, J., & Frahm, J. (2018). Central sulcus depth profiles of a large human cohort: Incidence of a divided central sulcus and description of the pli de passage fronto-parietal moyen Annual Meeting Society for Neuroscience San Diego, USA.

Schweizer, R., Helms, G., & Frahm, J. (2014). Revisiting a historic human brain with magnetic resonance imaging – the first description of a divided central sulcus [Original Research]. Frontiers in Neuroanatomy, 8. 10.3389/fnana.2014.00035

Schweizer, R., Müllen, A., & Frahm, J. (2019). Location and Height of the Central Sulcus Pli-de-Passage Fronto-Pariétal Moyen in a Large Cohort OHBM Annual Meeting, Rom, Italy.

Schweizer, R., Stropel, J., Grätz, H., & Frahm, J. (2018). Central Sulcus depth-profiles from structural MRI in humans: divided central sulci and pli de passage fronto-parietal moyen FENS Forum 2018, Berlin, Germany.

Schweizer, R., Wittmann, A., & Frahm, J. (2014). A rare anatomical variation newly identifies the brains of C.F. Gauss and C.H. Fuchs in a collection at the University of Göttingen. Brain, 137(4), e269. 10.1093/brain/awt296

Team, R. C. (2020). R: A Language and Environment for Statistical Computing. R Foundation for Statistical Computing.

Van Essen, D. C., Ugurbil, K., Auerbach, E., Barch, D., Behrens, T. E. J., Bucholz, R., Chang, A., Chen, L., Corbetta, M., Curtiss, S. W., Della Penna, S., Feinberg, D., Glasser, M. F., Harel, N., Heath, A. C., Larson-Prior, L., Marcus, D., Michalareas, G., Moeller, S., … Yacoub, E. (2012). The Human Connectome Project: A data acquisition perspective. NeuroImage, 62(4), 2222–2231. 10.1016/j.neuroimage.2012.02.018

von Monakow, C. (1905). *Gehirnpathologie* (zweite Auflage ed.). Alfred Hölder, K. U. K. Hof-und Universitäts-Buchhändler.

Wagner, R. (1860). Über die typischen Verschiedenheiten der Windungen der Hemisphären und über die Lehre vom Hirngewicht, mit besondrer Rücksicht auf die Hirnbildung intelligenter Männer. In Vorstudien zu einer wissenschaftlichen Morphologie und Physiologie des menschlichen Gehirns als Seelenorgan. Verlag der Dieterichschen Buchhandlung

Wagner, R. (1862). Über den Hirnbau der Mikrocephalen mit vergleichender Rücksicht auf den Bau des Gehirns der normalen Menschen und der Quadrumanen. In Vorstudien zu einer wissenschaftlichen Morphologie und Physiologie des menschlichen Gehirns als Seelenorgan. Verlag der Dietrichschen Buchhandlung.

White, T., Andreasen, N. C., & Nopoulos, P. (2002). Brain Volumes and Surface Morphology in Monozygotic Twins. Cerebral Cortex, 12(5), 486–493. 10.1093/cercor/12.5.486

